# The fast and superprocessive KIF1A predominately resides in a vulnerable one-head-bound state during its chemomechanical cycle

**DOI:** 10.1101/2020.06.28.176669

**Authors:** Taylor M. Zaniewski, Allison M. Gicking, John Fricks, William O. Hancock

## Abstract

Kinesin-3 are the fastest and most processive motors of the three neuronal transport kinesin families, yet the sequence of states and rates of kinetic transitions that comprise the chemomechanical cycle are poorly understood. We used stopped-flow fluorescence spectroscopy and single-molecule motility assays to delineate the chemomechanical cycle of the kinesin-3, KIF1A. Our bacterially expressed KIF1A construct, dimerized via a kinesin-1 coiled-coil, exhibits fast velocity and superprocessivity behavior similar to wild-type KIF1A. We established that the KIF1A forward step is triggered by hydrolysis of ATP and not by ATP binding, meaning that KIF1A follows the same chemomechanical cycle as established for kinesin-1 and-2. The ATP-triggered half-site release rate of KIF1A was similar to the stepping rate, indicating that during stepping, rear-head detachment is an order of magnitude faster than in kinesin-1 and kinesin-2. Thus, KIF1A spends the majority of its hydrolysis cycle in a one-head-bound state. Both the ADP off-rate and the ATP on-rate at physiological ATP concentration were fast, eliminating these steps as possible rate limiting transitions. Based on the measured run length and the relatively slow off-rate in ADP, we conclude that attachment of the tethered head is the rate limiting transition in the KIF1A stepping cycle. The fast speed, superprocessivity and load sensitivity of KIF1A can be explained by a fast rear head detachment rate, a rate-limiting step of tethered head attachment that follows ATP hydrolysis, and a relatively strong electrostatic interaction with the microtubule in the weakly-bound post-hydrolysis state.

## INTRODUCTION

The kinesin-3 motor protein KIF1A is a neuronal transport motor responsible for the anterograde transport of synaptic vesicle precursors and other vesicular cargo along microtubules (Mt).^1–4^ Mutations of KIF1A in humans can cause a range of afflictions known as KIF1A Associated Neurological Disorders (KAND) that include sensory and motor disabilities.^5–7^ In some cases, these disorders are caused by neuronal cell death and axon degeneration or specific mutations leading to the hyperactivation of KIF1A and an abundance of the correlative cargo at the synapse.^8^ However, in most cases the links between the motor dysfunction and the resulting disease are not clear.

The kinesin-3 family is one of the largest of the 14 subfamilies in the kinesin superfamily,^1,2,5,9–12^ and KIF1A is of particular interest due to a unique set of properties, including fast velocity,^13^ superprocessivity,^13–16^ low force resistance^17^ and the ability to move processively as both a monomer and dimer.^18–21^ The superprocessivity (long travel distance before detaching) of KIF1A has been explained by an electrostatic interaction between the positively charged loop-12 of KIF1A called the ‘K-loop’ and the negatively-charged C-terminal tail of tubulin.^15,17,18,22–26^ An adaptation that increases microtubule affinity would generally be expected to slow the velocity rather than speed it up,^27,28^ yet KIF1A steps 2.5-fold faster than kinesin-1. Furthermore, optical trapping studies and mixed motor assays have revealed that, despite the enhanced electrostatic association to the microtubule, kinesin-3 has a surprisingly low resistance to force and detaches under load.^8,15,17,29–31^ How these opposing traits are reconciled in the same motor have yet to be fully understood.

Interpreting the chemomechanical properties of KIF1A and how the motor is tuned for its specific cellular functions requires a more complete understanding of the KIF1A chemomechanical cycle. Specifically, it remains to be determined whether the fast speed and superprocessivity of KIF1A result simply from differences in specific rate constants in the hydrolysis cycle, or whether they result from the KIF1A cycle having a different sequence of chemomechanical states than kinesin-1. In the kinesin-1 chemomechanical cycle (Fig. 1), it has been established that, following initial binding and release of ADP (state 3), kinesin-1 waits for ATP binding with the tethered head in a rearward position.^27,32,33^ ATP binding to the bound head then repositions the tethered head forward, and ATP hydrolysis triggers full neck linker docking, which positions the tethered head near its next binding site. The forward step is completed by the tethered head binding the microtubule and releasing its bound ADP to generate a tight-binding state 7.^27,28,34–37^ The key transition that determines processivity in this model is the kinetic race out of state 5 – the race is won if the tethered head binds the next tubulin before the bound head detaches from the vulnerable ADP-Pi state. Therefore, processivity requires that the rate of tethered head attachment be considerably faster than the rate of bound head dissociation from the microtubule.

**Figure 1.**
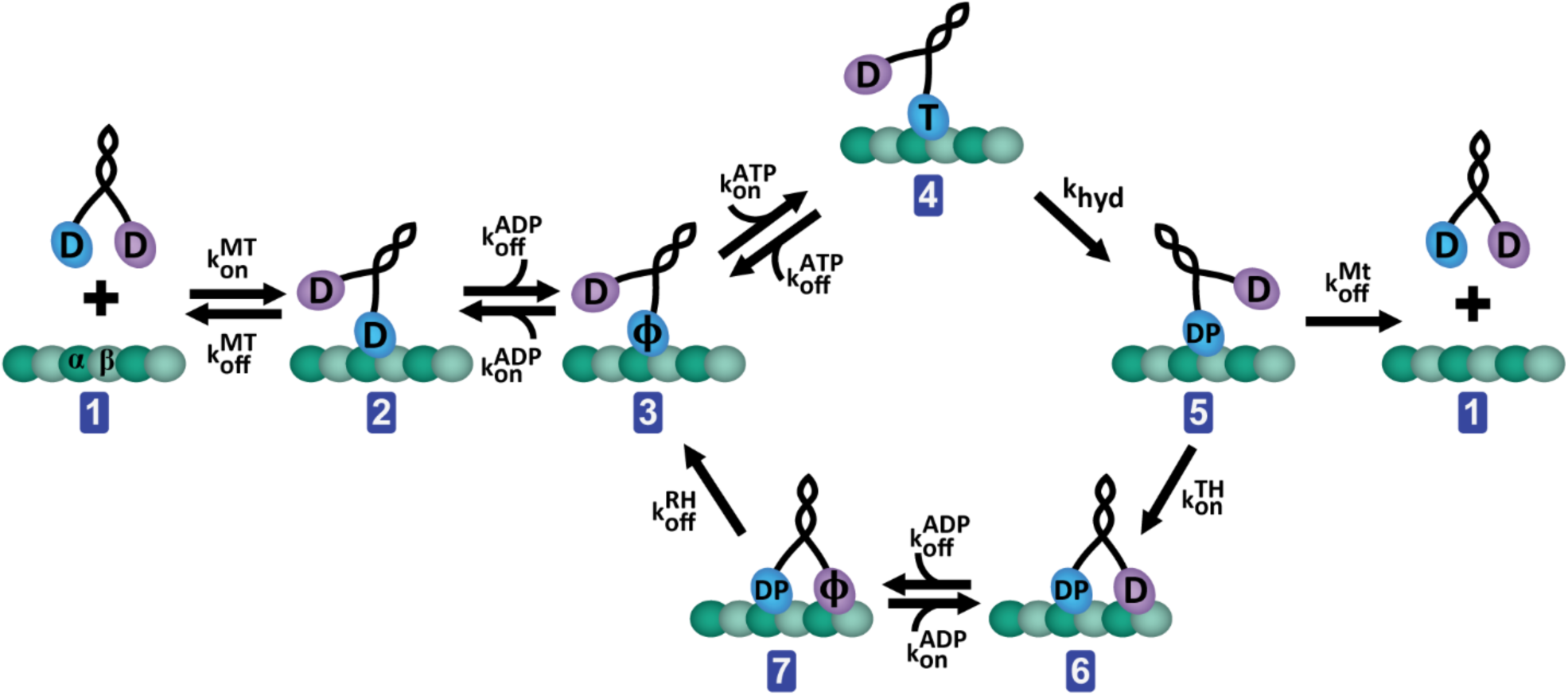
Canonical kinesin chemomechanical cycle. The motor protein begins with ADP bound to both motor domains in solution (state 1). Upon binding to the microtubule (state 2), one ADP is released, locking the motor in a strongly bound state, while the other ADP remains bound to the tethered head (state 3). ATP then binds to the bound head (state 4) and is hydrolyzed to ADP-Pi (state 5), triggering full neck linker docking, which positions the tethered head forward and puts the motor in a weakly-bound state.^27^ From this vulnerable state 5, the bound head can detach from the microtubule and terminate the processive run (state 1). More often, the tethered head binds to its next binding site (state 6) and ADP is released to generate a tightly-bound state 7 that completes the forward step. Detachment and Pi release by the rear head returns the motor to the ATP binding state (state 3).^27,28,34–37^ D=ADP; T=ATP; DP= ADP-P_i;_ ϕ=Apo.

Mapping this canonical kinesin-1 hydrolysis cycle to the characteristics of KIF1A, one or more transitions must be ∼2.5 times faster than kinesin-1 to account for the faster stepping rate, and the probability of dissociating per cycle must be ∼7-fold lower to account for its superprocessivity.^27,28^ Although there are several ways this may be achieved, one intriguing possibility is KIF1A bypassing specific transition states in the cycle. By removing the need for the forward step to be triggered by either ATP hydrolysis (removing state 5) or ATP binding (removing both states 4 and 5), KIF1A could theoretically both reduce the total number of sequential forward transitions in the cycle (gaining speed) and avoid the vulnerable one-head-bound ADP-Pi State 5 (enhancing processivity). The goal of the present work is to use single-molecule tracking and pre-steady-state kinetic analysis of dimeric KIF1A to define the sequence of states that make up the KIF1A chemomechanical cycle and quantify the transition rates between these states. By delineating the chemomechanical cycle of KIF1A, we provide a mechanistic explanation of the motor’s superprocessivity, high velocity and sensitivity to load.

## RESULTS

To form a stable dimer and allow for direct comparison of the properties of KIF1A to kinesin-1 and kinesin-2 constructs characterized previously^27,28,34^, we bacterially expressed a *Rattus norvegicus* KIF1A construct dimerized via the *Drosophila melanogaster* KHC neck coil (Fig. 2A and 2B). Two different lengths of the kinesin-1 neck coil were used in these KIF1A constructs for distinct purposes. For biochemical assays, we used KIF1A-406, which includes 61 residues from the kinesin-1 (*Dm*KHC) neck coil added after the native KIF1A head and neck linker. For microscopy, we used KIF1A-560-GFP, which includes 216 residues from kinesin-1 that include the neck coil and coil-1, followed by a C-terminal GFP. All experiments were carried out in 80 mM PIPES buffer (BRB80) to ensure physiologically relevant ionic strength of the solution.

**Figure 2.**
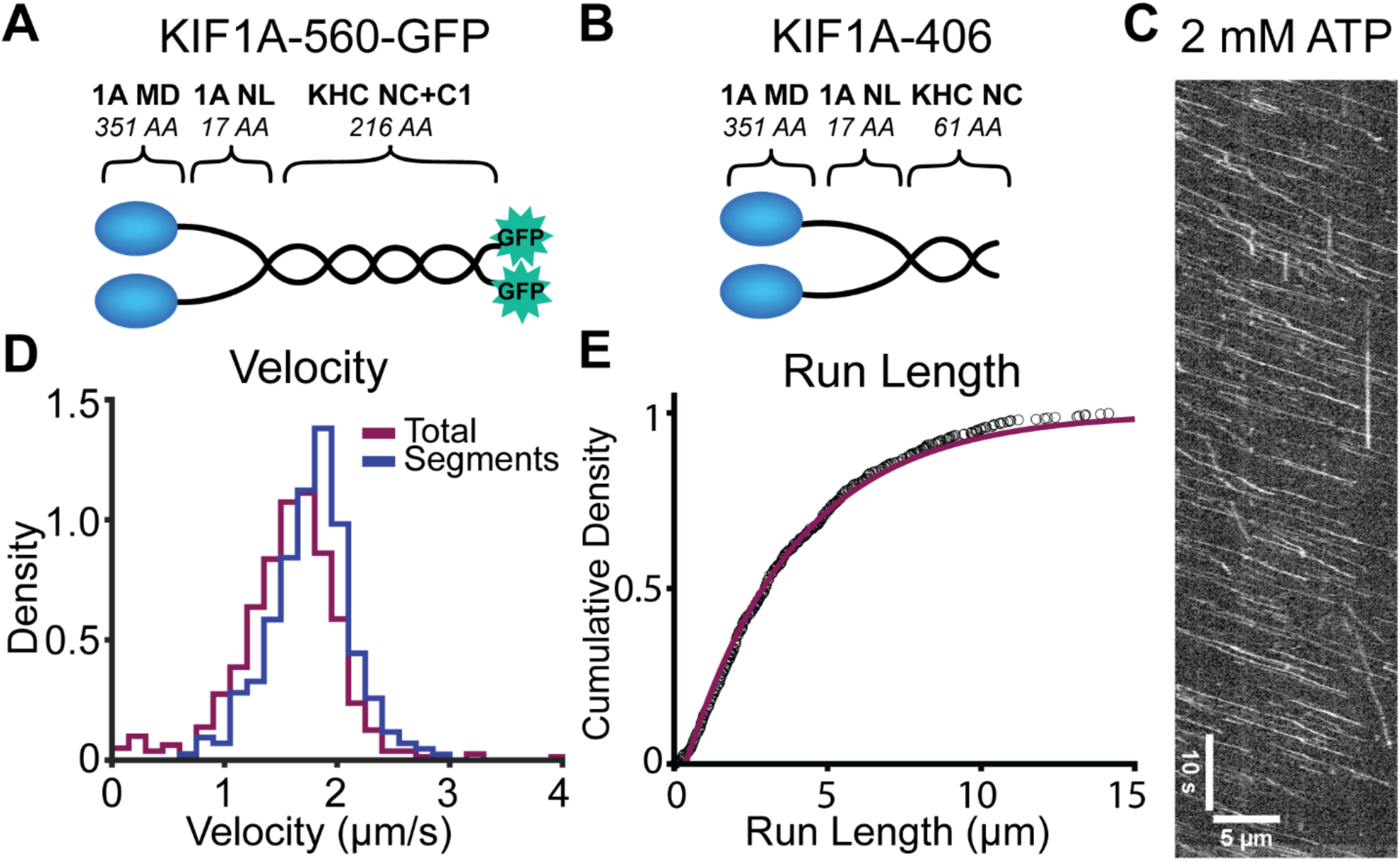
Bacterially expressed KIF1A dimer is fast and superprocessive. **A**, Diagram of KIF1A-560-GFP construct used in single-molecule assays. **B**, Diagram of KIF1A-406 construct used in biochemical assays. (Diagrams in A and B are not to scale). **C**, Kymograph of KIF1A-560-GFP motility in 2 mM ATP at 10 fps. **D**, Histogram of velocities determined from measuring the total trace (including pauses), and the linear regions of traces (excluding pauses). Mean velocities were 1.56 ± 0.5 μm/s (mean ± SD, N = 534) for total traces and 1.77 ± 0.4 μm/s (mean ± SD, N = 285) for linear regions. **E**, Single-molecule run length of 3.6 ± 0.04 μm (mean ± 95% confidence, N=534) was determined by cumulative density fit to the run lengths above 0.4 μm. Statistical analysis of the traces terminated by microtubule length gives an estimated total run length of 5.6 ± 0.4 μm (see Methods). *Abbreviations*, 1A, KIF1A; MD, motor domain; NL, neck linker; KHC, kinesin heavy chain; NC, neck coil; C1, coil-1; GFP, green fluorescent protein.

### KIF1A is fast and superprocessive

Using single-molecule TIRF microscopy at 2 mM ATP, we measured the velocity of KIF1A-560-GFP from kymograph evaluation by the two following methods: 1) linear segments of uninterrupted stepping (excluding pauses) and 2) total runs (including pauses) (Fig. 2C). We determined a velocity of 1.77 ±0.4 μm/s (mean ± SD, N = 285), when pauses are excluded, and 1.56 ± 0.5 μm/s (mean ± SD, N = 534) for entire runs (Fig. 2D). Assuming an 8-nm step size, these velocities translate to stepping rates of 220 ± 50 s^−1^ and 195 ± 63 s^−1^, respectively. In addition to velocity, we also determined a run length of 3.6 ±0.04 μm (mean ± 95% confidence interval) (Fig. 2E). This run length is an underestimate due to the finite microtubule lengths in the assay, a common limitation seen in KIF1A studies.^13–15^ Therefore, we developed a statistical model that accounts for runs that terminate prematurely due to motors reaching the end of the microtubule (see Methods). Using this correction, we estimate a true average run length of 5.6 ± 0.4 μm (mean ± SD). Considering this estimated run length and the velocity over the total trace, we determined a mean run time of 3.6 ± 1.2 s, corresponding to a motor off-rate of 0.28 ± 0.09 s^−1^. The velocity and run length determined here for our bacterially expressed KIF1A dimer are consistent with the fast velocity and superprocessivity reported in previous studies.^14,15^

### ATP Binding and Hydrolysis are both required for fast forward stepping of KIF1A

The features of the KIF1A chemomechanical cycle that underlie the fast velocity and superprocessivity are not known. Based on the similarities in structure and cellular function to kinesin-1, a logical hypothesis is that kinesin-3 follows the same chemomechanical model that has been delineated for kinesin-1. However, this has never been experimentally confirmed.^27,28,36,38,39^ Furthermore, it is possible that, rather than resulting from quantitative differences in rate constants between the kinesin families, the faster speed and enhanced processivity of KIF1A may result from qualitative differences in the sequence of states that make up the chemomechanical cycle. One candidate is the state that triggers the forward step. Recent experiments demonstrated that instead it is ATP hydrolysis, rather than ATP binding alone, that triggers the forward step in kinesin-1.^36,38^ In principle, reducing the time the motor spends in the 1HB state, either by removing the need for an ATP-binding or ATP-hydrolysis step-trigger, could increase the overall stepping rate by speeding a key process in the cycle, and increase the processivity of the motor by reducing the probability of the motor dissociating before completing its forward step. To this end, we designed a series of experiments to ask whether the sequence of biochemical states that triggers the forward step of KIF1A matches those of kinesin-1 and -2. As shown in Fig. 3A, following detachment of the rear head from the rear binding site, there are three potential events that could trigger the KIF1A forward step: 1) the forward step could occur spontaneously while the bound head is in the Apo state; 2) ATP binding to the bound head could trigger the forward step; or 3) ATP hydrolysis by the bound head could trigger the forward step.

**Figure 3.**
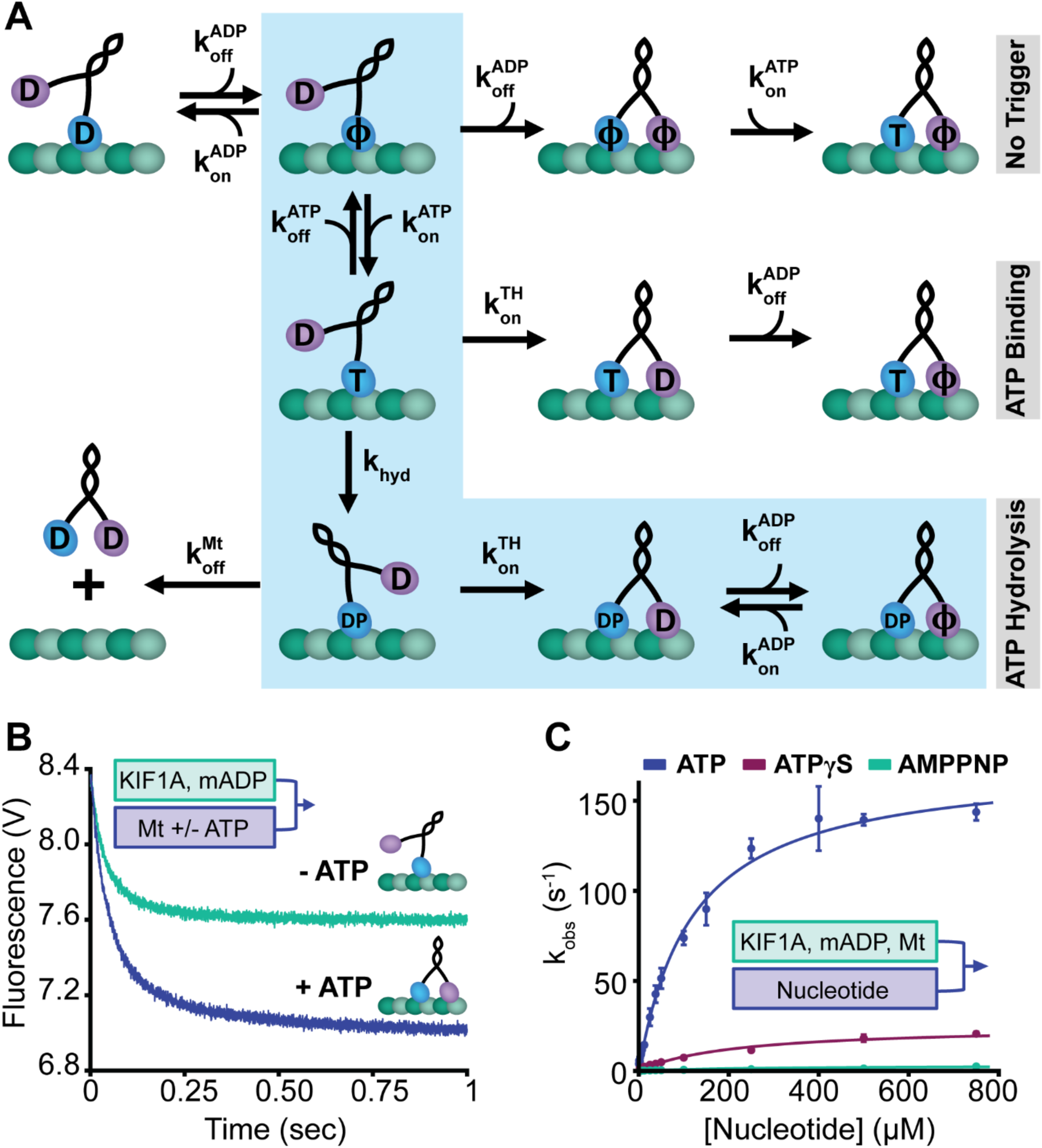
The forward step of KIF1A is triggered by ATP Hydrolysis. **A**, Diagram of three proposed models for the stepping trigger in the KIF1A chemomechanical cycle. In the No Trigger model, the tethered head steps independent of the nucleotide state of the bound head. In the ATP Binding model, ATP binding to the bound head triggers the forward step by the tethered head. In the ATP Hydrolysis-triggered model, highlighted by light blue shading, ATP hydrolysis is required for forward stepping by the tethered head. D=ADP; T=ATP; DP= ADP-P_i;_ ϕ=Apo. **B**, KIF1A half-site reactivity experiment. 150 nM of mADP-labelled KIF1A was flushed against a solution of 1 μM microtubules either with or without 1 mM ATP (all final chamber concentrations). Amplitudes of the traces were 0.7 V in the absence of nucleotide and 1.2 V in the presence of ATP. **C**, Nucleotide-triggered Half-Site Release Assay. 150 nM of mADP-exchanged KIF1A and 3 μM microtubules were flushed against varied concentrations of the ATP, ATPγS, or AMPPNP (all final chamber concentrations). Fitting with a hyperbola gave maximal rates of 172 ± 10 s^−1^, 25 ± 6 s^−1^, and 0.44 ± 0.03 s^−1^ for ATP, ATPγS, and AMPPNP, respectively (fit ± 95% confidence interval). The corresponding K_0.5_ values were 119 ± 21 μM, 215 ± 110 μM, and 22 ± 7 μM for ATP, ATPγS, and AMPPNP, respectively (fit ± 95% CI).

To test whether the forward step can occur spontaneously, we asked if, upon binding to the microtubule in the absence of free nucleotide, the motor releases one or both bound ADP (see Methods for details).^40^ If no trigger is required for the forward step, then the motor should release both ADP upon nucleotide binding, whereas if a trigger is required for the forward step, then only one ADP will be released. To measure the release of the nucleotide from the motor head domain, we used mant-ADP whose fluorescence is enhanced upon motor binding. Thus, mADP dissociation from the motor can be monitored by a decrease in mADP fluorescence. In the control experiment, KIF1A in mADP was flushed against microtubules and 1 mM ATP, which triggers stepping and rapid release of both bound mADP (Fig. 3B, blue trace). In the absence of ATP, however, the fluorescence only decreased by half, indicating that KIF1A released only half of its nucleotide upon microtubule binding (Fig. 3B, green trace). Thus, a trigger in the form of ATP binding or ATP hydrolysis by the bound head is necessary to catalyze the forward step of KIF1A. This result nullifies the first potential pathway in Fig. 3A.

To determine whether ATP binding alone is sufficient to trigger the forward step or if hydrolysis is necessary, we used a nucleotide-triggered half-site release assay first used by Ma and Taylor.^41^ In this experiment, motors and microtubules are combined in the absence of free nucleotide to produce a 1HB ATP waiting state with mADP in the tethered head. Different concentrations of ATP or ATP analogs are then flushed against this complex and the rate of mADP release from the tethered head is measured. ATP triggered a maximal half-site release rate of 172 ± 10 s^−1^ (fit ± 95% CI) (Fig. 3C, blue trace), which is similar to the motor stepping rate. If only ATP binding is required to trigger the step, then the slowly hydrolyzed ATP analog ATPγS or the nonhydrolyzable ATP analog AMPPNP should also trigger half-site release at a similar rate. Instead, ATPγS triggered half-site release of only 24.9 ± 6.4 s^−1^ (fit ± 95% CI) (Fig. 3C, red trace) and AMPPNP triggered a maximal half-site release rate of only 0.44 ± 0.03 s^−1^ (fit ± 95% CI) (Fig. 3C, green trace). These rates are both significantly slower than either the ATP-triggered half-site release rate or the stepping rate, thus nullifying our second potential pathway in Fig. 3A. In a control experiment, the KIF1A single-molecule velocity in 1 mM ATPγS was 180 ± 0.2 nm/s (mean ± SD, data not shown), corresponding to 23 steps/s and indicating that the elevated half-site release in ATPγS compared to AMPPNP likely results from ATP hydrolysis. Thus, we conclude that, during the normal stepping cycle, ATP-hydrolysis is required to trigger the forward step, which is completed by the forward head releasing ADP to generate a tightly-bound state (shaded pathway in Fig. 3A). The observation that both ATP binding and hydrolysis are required for the forward step indicates that KIF1A follows a similar hydrolysis cycle to kinesin-1 and -2,^27,34^ and therefore the enhanced motility must result from quantitative differences in transition rates between each state.

### Transition rates in the KIF1A chemomechanical cycle

Having defined the sequence of the states in the KIF1A chemomechanical cycle, we then measured the kinetic rates of each of the transitions KIF1A undergoes upon interaction with the microtubule. Preceding the stepping cycle, the motor protein must first land on the microtubule. Therefore, to gain insight into the KIF1A-microtubule affinity, we measured the microtubule on-rate in solution (step 1→2 in Fig. 1). By flushing motors against varying concentrations of microtubules, we monitored mADP release from the motor upon microtubule binding.^34^ When mADP-bound motors are flushed against low concentrations of microtubules, the microtubule binding step is rate limiting, enabling determination of the first-order on-rate for microtubule binding. From this assay, we calculated a 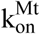 of 17 ± 4 μM^−1^s^−1^. (Fig. 4A, fit ± 95 % CI).

**Figure 4.**
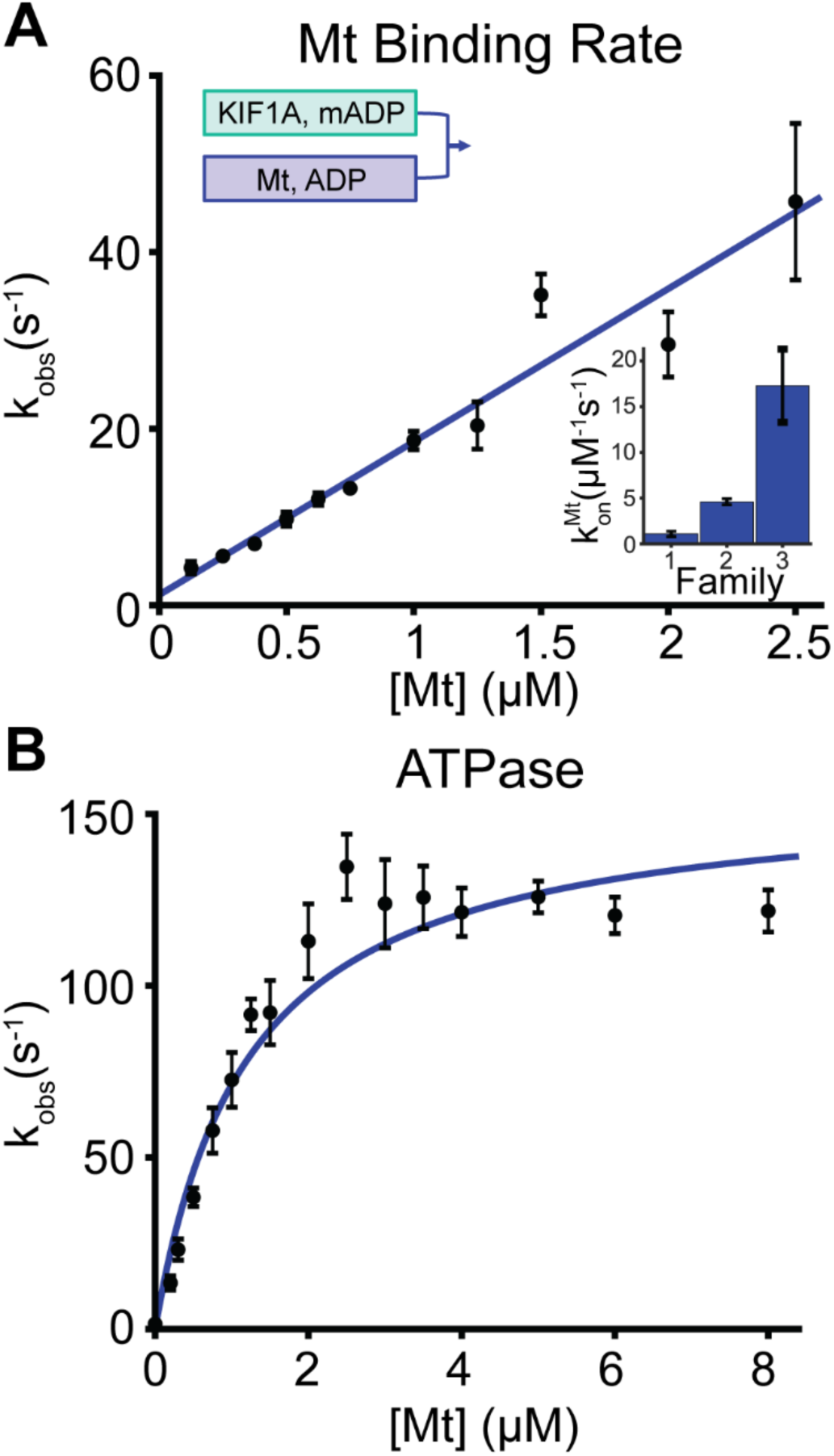
KIF1A ATPase and Microtubule on-rate. **A**, KIF1A microtubule on-rate, measured by mADP release by the motor upon binding to the microtubule. A linear fit to the observed rates as a function of microtubule concentration gave a bimolecular on-rate, 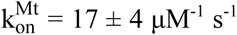 (fit ± 95% CI; N=3 trials per point with N=5-7 shots per trial; error bars are SEM). **Inset:** Comparing microtubule binding rates in BRB80 for kinesin-1, -2 and -3.^34,42^ **B**, Microtubule-stimulated ATPase of KIF1A. A Michaelis-Menten fit weighted by the inverse of SEM of the points (N=6 trials per point), gave a k_cat_ of 115 ± 16 s^−1^ and a K_m_ of 1.2 ± 0.5 μM (fit ± 95% confidence interval).

Notably, this rate is approximately 15-fold faster than the corresponding rate for kinesin-1^42^ (Table 2) and is consistent with fast KIF1A single-molecule landing rates observed previously.^15^ The second question we addressed was whether ATP hydrolysis is tightly coupled to motor stepping; if the motor undergoes futile hydrolysis cycles during stepping, then the Fig. 1 model will have to be modified to explain KIF1A. To measure the ATP hydrolysis cycle rate, we used an enzyme-coupled assay to measure the KIF1A ATPase at varying microtubule concentrations. Fitting with the Michaelis-Menten equation, we measured a k_cat_ of 115 ± 16 s^−1^ and a K_m_ of 1.2 ± 0.5 μM (Fig. 4B, fit ± 95% CI). This k_cat_ is lower than the total stepping rate of 195 ± 63 s^−1^, determined from single-molecule velocity including pauses (Fig. 2D), arguing against the motor undergoing any futile cycles of ATP hydrolysis under no load. The k_cat_ calculated here may be underestimated since the active motor concentration determined by microtubule pelleting assay in AMPPNP (see Methods) may be an overestimation due to inactive motors that irreversibly bind. Thus, because our transient kinetics investigations are generally studying only one motor step, we choose to use the uninterrupted stepping rate at 25°C of 220 ± 50 s^−1^ (Fig. 2D) as the best estimate of the overall KIF1A chemomechanical cycle rate.

**Table 1.** Rates and state durations of the KIF1A chemomechanical cycle.

| Parameter | Notation | Experimental | Duration | Source |
| --- | --- | --- | --- | --- |
| Velocity (with pauses) | $Vel_a$ | $1.56 \pm 0.5$ $\mu\text{m/s}$ | | Fig 2D |
| Velocity (without pauses) | $Vel_b$ | $1.77 \pm 0.4$ $\mu\text{m/s}$ | | Fig 2D |
| Run Length (Measured) | $RL_a$ | $3.6 \pm 0.04$ $\mu\text{m}$ | | Fig 2E |
| Run Length (Corrected) | $RL_b$ | $5.6 \pm 0.4$ $\mu\text{m}$ | | Eq. 12 |
| Step Number | Steps | $700 \pm 50$ steps | | $RL_b / 8 \text{ nm}$ |
| Stepping rate | $k_{\text{step}}$ | $220 \pm 50$ $\text{s}^{-1}$ | $4.5 \pm 1.0$ ms | $Vel_b / 8 \text{ nm}$ |
| Mt off-rate in ATP | $k_{\text{off}}^{\text{Mt}}$ | $0.28 \pm 0.09$ $\text{s}^{-1}$ | $3.6 \pm 1.2$ s | $Vel_a / RL_b$ |
| Mt off-rate in ADP | $k_{\text{off}}^{\text{Mt}}$ | $0.27 \pm 0.11$ $\text{s}^{-1}$ | $3.72 \pm 0.03$ s | Fig 6D |
| Half-site release rate | $k_{\text{max}}^{\text{HS}}$ | $172 \pm 10$ $\text{s}^{-1}$ | $5.8 \pm 0.4$ ms | Fig 3C |
| ATP for half-max release | $K_{0.5}^{\text{HS}}$ | $119 \pm 21$ $\mu\text{M ATP}$ | | Fig 3C |
| ATP on-rate (lower limit) | $k_{\text{on}}^{\text{ATP}}$ | $\geq 1.4 \pm 0.3$ $\mu\text{M}^{-1} \text{s}^{-1}$ | $\leq 0.7 \pm 0.15$ ms | Fig 3C |
| Mt on-rate | $k_{\text{on}}^{\text{Mt}}$ | $17 \pm 4$ $\mu\text{M}^{-1} \text{s}^{-1}$ | | Fig 4A |
| ATPase Cycle Rate | $k_{\text{cat}}$ | $115 \pm 16$ $\text{s}^{-1}$ | $8.7 \pm 1.2$ ms | Fig 4B |
| Michaelis-Menten constant | $K_M$ | $1.2 \pm 0.5$ $\mu\text{M Mt}$ | | Fig 4B |
| Strained mADP off-rate | $k_{\text{off}}^{\text{mADP}}$ | $616 \pm 86$ $\text{s}^{-1}$ | $1.6 \pm 0.2$ ms | Fig 5A |
| Unstrained mADP off-rate | $k_{\text{off}}^{\text{mADP}}$ | $354 \pm 78$ $\text{s}^{-1}$ | $2.8 \pm 0.6$ ms | Fig 5B |
| Unstrained mADP on-rate | $k_{\text{on}}^{\text{mADP}}$ | $29 \pm 15$ $\mu\text{M}^{-1} \text{s}^{-1}$ | | Fig 5B |
| Solution ADP off-rate | $k_{\text{off}}^{\text{ADP}}$ | $0.26 \pm 0.001$ $\text{s}^{-1}$ | $3.8 \pm 0.02$ s | Fig 5C |
| Solution mADP off-rate | $k_{\text{off}}^{\text{mADP}}$ | $0.27 \pm 0.005$ $\text{s}^{-1}$ | $3.7 \pm 0.07$ s | Fig 5D |
| Tethered-head on-rate | $k_{\text{on}}^{\text{TH}}$ | $189 \pm 78$ $\text{s}^{-1}$ | $5.3 \pm 2$ ms | Eq 1 |
| Hydrolysis rate | $k_{\text{hyd}}$ | Fast* $\text{s}^{-1}$ | Fast* ms | Eq 3 |
| Rear-head detachment rate | $k_{\text{off}}^{\text{RH}}$ | Fast* $\text{s}^{-1}$ | Fast* ms | Eq 4 |

\*Fast refers to rates that are above our detection limit.

**Table 2.** Comparing kinetic parameters for the kinesin families 1, 2 and 3. *Some kinesin-2 values are approximate due to the motor’s higher affinity for mADP than unlabeled-ADP, therefore, no error is reported. References: **a**, Feng *et al*. 2018^42^; **b**, Chen *et al*. 2015^34^; **c**, Mickolajczyk and Hancock 2017^28^; **d**, Mickolajczyk *et al*. 2015^27^; **e**, This Study. Calculations and errors are propagated from reported values in reference papers.

|  | Kinesin-1 |  |  | Kinesin-2 |  |  | Kinesin-3 |  |  |
| --- | --- | --- | --- | --- | --- | --- | --- | --- | --- |
| Parameter | Experimental | Units | Ref | Experimental | Units | Ref | Experimental | Units | Ref |
| $k_{on}^{Mt}$ | $1.1 \pm 0.05$ | $\mu M^{-1} s^{-1}$ | a | $4.6 \pm 0.9$ | $\mu M^{-1} s^{-1}$ | b | $17 \pm 4$ | $\mu M^{-1} s^{-1}$ | e |
| $k_{on}^{ATP}$ | $> 1.2 \pm 0.3$ | $\mu M^{-1} s^{-1}$ | d | 18.0 | $\mu M^{-1} s^{-1}$ | b | $> 1.4 \pm 0.3$ | $\mu M^{-1} s^{-1}$ | e |
| $k_{hyd}$ | $281 \pm 215$ | $s^{-1}$ | c | $478 \pm 489$ | $s^{-1}$ | b | n.d. | $s^{-1}$ | e |
| $k_{on}^{TH}$ | $216 \pm 22$ | $s^{-1}$ | c | $117 \pm 14$ | $s^{-1}$ | b | $189 \pm 78$ | $s^{-1}$ | e |
| $k_{off}^{ADP}$ | $367 \pm 4$ | $s^{-1}$ | d | 390* | $s^{-1}$ | b | $615 \pm 86$ | $s^{-1}$ | e |
| $k_{off}^{RH}$ | $154 \pm 17$ | $s^{-1}$ | d | $89 \pm 25$ | $s^{-1}$ | b | n.d. | $s^{-1}$ | e |
| $k_{cat}$ | $67 \pm 11$ | $s^{-1}$ | c | $42 \pm 4$ | $s^{-1}$ | b | $115 \pm 16$ | $s^{-1}$ | e |
| $k_{max}^{HS}$ | $112 \pm 9$ | $s^{-1}$ | d | $\geq 47^*$ | $s^{-1}$ | b | $172 \pm 10$ | $s^{-1}$ | e |
| Vel | $533 \pm 3$ | nm/s | d | $400 \pm 40$ | nm/s | c | $1560 \pm 500$ | nm/s | e |
| RL | $860 \pm 20$ | nm | c | $550 \pm 70$ | nm | c | $5600 \pm 400$ | nm | e |
| $k_{off}^{Mt \text{ in ATP}}$ | $0.81 \pm 0.14$ | $s^{-1}$ | c | $0.73 \pm 0.12$ | $s^{-1}$ | c | $0.28 \pm 0.09$ | $s^{-1}$ | e |
| $k_{off}^{Mt \text{ in ADP}}$ | $2.0 \pm 0.2$ | $s^{-1}$ | c | $2.3 \pm 0.2$ | $s^{-1}$ | c | $0.27 \pm 0.11$ | $s^{-1}$ | e |
| # Steps | $108 \pm 3$ | steps | c | $69 \pm 9$ | steps | c | $700 \pm 50$ | steps | e |
| $k_{step}$ | $65 \pm 0.4$ | $s^{-1}$ | d | $50 \pm 5$ | $s^{-1}$ | c | $220 \pm 50$ | $s^{-1}$ | e |

To identify the rate limiting step in the KIF1A cycle, we designed experiments to measure the rates of the specific transitions within the cycle and compared them to the overall stepping rate. Possible transitions that could determine the overall KIF1A cycle rate (Fig. 1) include: 1) ATP binding 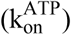, 2) ATP hydrolysis (k_hyd_), 3) tethered-head attachment to the next tubulin 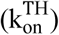, 4) ADP release by the tethered head 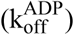, and 5) rear-head detachment 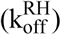.

### ATP binding and ADP release are not rate-limiting

The first portion of the stepping cycle that can be excluded as a possible rate-limiting step is the ATP on-rate (state 3 → 4 in Fig. 1). This can be shown by the observation that in the ATP-triggered half-site release experiment in Fig. 3C, the maximal rate was 172 s^−1^, and the half-maximal rate was achieved at an ATP concentration of 119 μM. Thus, at 1 mM ATP the curve has reached a plateau indicating that ATP binding is not rate limiting. Going further, the half-max (K_0.5_) can be used to estimate a lower limit for ATP binding, as follows: If ATP binding were irreversible and the reaction is treated as a sequence of ATP binding followed by the remainder of steps, then it follows that at the ATP concentration that produces half-maximal release, half of the time is taken by ATP binding. At saturating ATP (where ATP binding is very fast), the release rate is 172 s^−1^, meaning that at the K_0.5_ of 119 μM ATP, the binding rate of ATP is 172 s^−1^ (followed by the remainder of the steps at 172 s^−1^). This K_0.5_ corresponds to a second-order on-rate for ATP binding of 172 s^−1^ / 119 μM = 1.4 μM^−1^s^−1^, which at 1 mM ATP corresponds to a rate of 1400 s^−1^, much faster than the 220 s^−1^ stepping rate. Also, if ATP binding is reversible, which is likely the case, then the on-rate would need to be even faster. We therefore conclude that at physiological ATP concentrations, ATP binding is far from rate limiting in the KIF1A hydrolysis cycle.

The second step we were able to rule out as rate-limiting is ADP release (state 6→7 in Fig. 1). To do this we measured the rate of nucleotide exchange assays in the strained two-head-bound (2HB) state. We generated a 2HB state by incubating KIF1A with microtubules in the presence of AMPPNP, which results in the rear head trapping the nonhydrolyzable nucleotide and the front head being trapped in a tight-binding apo state.^27,34,43,44^ Flushing this complex against mADP results in reversible nucleotide binding to the leading head. From this experiment, we determined an ADP off-rate of 616 ± 86 s^−1^ (mean ± SD) from the strained leading head (Fig 5A). As this measurement is near the limit of the instrument’s capabilities, there was not a clear increase in the observed rate with increasing mADP concentrations and thus, our estimate represents an average across nucleotide concentrations. To measure microtubule-stimulated ADP off-rate in a different way, we measured the rate of mADP exchange when the motor is bound to the microtubule in the one-head-bound state. As shown in the half-site release experiment (Fig. 3C), incubating KIF1A with microtubules in the absence of added nucleotide results in release of one ADP and formation of a 1HB complex. By flushing this complex against different concentrations of mADP, we measured an unstrained ADP on-rate of 29 ± 15 μM^−1^ s^−1^ and unstrained ADP off-rate of 354 ± 78 s^−1^ (Fig 5B, fit ± 95% CI). Although this unstrained ADP off-rate is likely less relevant to the normal stepping cycle than the strained rate, it is still faster than the 220 s^−1^ overall stepping rate. Finally, to rule out the possibility that mADP off-rates are not representative of unlabeled ADP, we measured ADP off-rates from KIF1A in the absence of microtubules. From these assays (see Methods for details), the unlabeled-ADP off-rate of 0.26 ± 0.005 s^−1^ (fit ± 95% CI, Fig 5C) was in good agreement with the mADP off-rate of 0.27 ± 0.001 s^−1^ (fit ± 95% CI, Fig 5D). Notably, these solution off-rates were roughly 20-fold faster than the corresponding ADP off-rate for kinesin-1, which is approximately 0.01 s^−1^.^45^ Although this off-rate in solution does not play a part in the normal ATP-stimulated chemomechanical cycle on the microtubule, it is indicative of differences in the nucleotide binding affinity that may relate to the fast KIF1A stepping speed. In summary, the ∼600 s^−1^ strained mADP off-rate, the ∼350 s^−1^ unstrained mADP off-rate, and the similarity in solution off-rates for ADP and mADP argue strongly that ADP release is not the rate limiting step in the overall stepping cycle of KIF1A.

**Figure 5.**
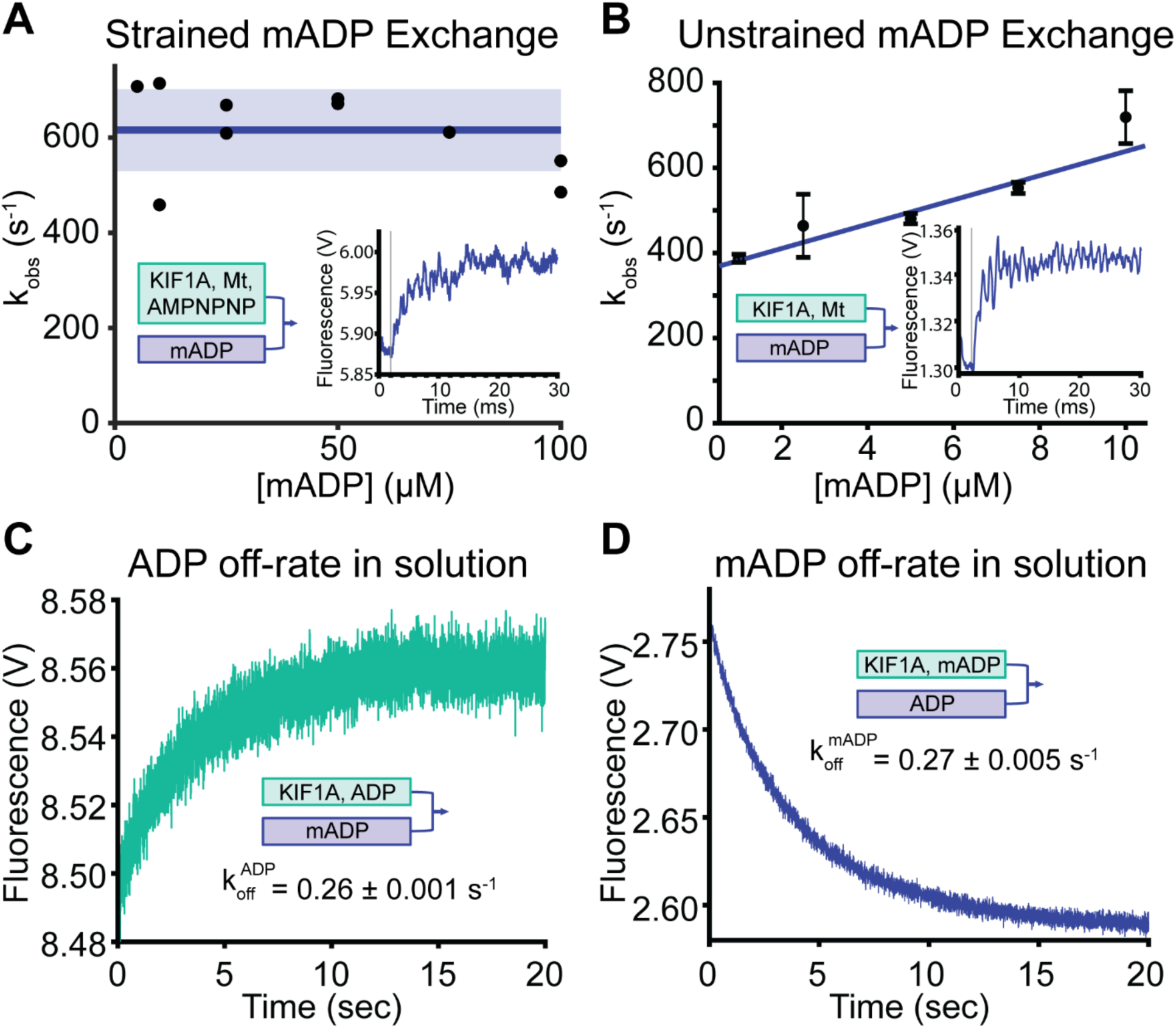
KIF1A has a high ADP off-rate. **A**, Exchange rate of mADP in the front head of KIF1A when the motor is in the 2HB state. 1 μM KIF1A, 5 μM microtubules and 50 μM AMPPNP were combined and flushed against varied [mADP] (all final chamber concentrations). Solid blue line at 616 s^−1^ indicates the mean rate across all [mADP]. (Shaded region is ± SD, N=2 trials) **Inset**, raw trace of stopped flow results at final [mADP] = 100 μM, N=6 traces averaged. Grey line at 2 ms indicates start of fit. **B**, Exchange rate of mADP in the bound head of KIF1A when the motor is in the 1HB state. 0.5 μM mADP-exchanged KIF1A dimers, 2.5 μM microtubules and 0.25 μM mADP were combined and flushed against varied [mADP] (all final chamber concentrations). Linear fit using 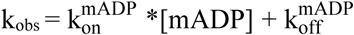gives 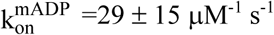 and 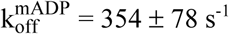 (N=3 trials per point, fit ± 95% CI, error bars are SEM). **Inset**, raw trace of stopped-flow results at 10 μM mADP, N=6 traces averaged. **C**, Time course of ADP dissociation from KIF1A in the absence of microtubules, triggered by flushing 0.15 μM motors in 0.25 μM unlabeled ADP against 5 μM mADP (all final chamber concentrations). An exponential fit, which is governed by the off-rate of unlabeled ADP, gives 0.26 ± 0.001 s^−1^. (Fit ± 95% CI, N=5-7 traces averaged). **D**, Time course of mADP dissociation from KIF1A in the absence of microtubules, triggered by flushing 0.15 μM motors and 0.25 μM of mADP against 1 mM unlabeled ADP (all final chamber concentrations). Exponential fit gives 0.27 ± 0.005 s^−1^ (Fit ± 95% CI, N=5-7 traces averaged).

### Rear-head detachment is fast

In the kinesin-1 and kinesin-2 chemomechanical cycles, rear-head detachment is at least partially rate-limting.^27,34,42^ This rate (State 7→3 in Fig. 1) can be calculated from the difference between the duration (inverse of the rate constant) observed in the ATP-triggered half-site release assay (States 3-7 in Fig. 1) and the total step duration (inverse of the stepping rate). Based on previous work, kinesin-1 has a step duration of 15.4 ms and spends 6.5 ms transitioning from the 2HB to 1HB state during rear-head detachment (Table 2).^27^ Similarly, rear-head detachment in kinesin-2 (11.2 ms) makes up 50% of the total cycle time (22.4 ms) (Table 2).^34^ To determine whether kinesin-3 follows this same trend, we compared the ATP-triggered half-site release rate (Fig. 3C) to the stepping rate. The pause-free stepping rate of 220 s^−1^ (Fig. 2D) converts to a step duration of 4.5 ± 1.0 ms. The maximal ATP-triggered half-site release rate of 172 s^−1^ (Fig. 3C) corresponds to a duration of 5.8 ± 0.4 ms. The similarity of these durations means that 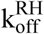 is faster than we can measure and that the rear-head detachment rate is not the rate-limiting step in the KIF1A hydrolysis cycle. Thus, the motor spends only a small fraction of its hydrolysis cycle in a 2HB state.

### Tethered-head attachment is rate-limiting

Since we have excluded 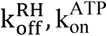, and 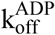 as potential rate limiting steps of the cycle, we are left with the rate-limiting step being either ATP hydrolysis (k_hyd_) or tethered head attachment 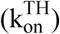. Measuring k_hyd_ generally requires quenched flow approaches, which are technically challenging for such a fast motor. However, because processivity can be considered as a kinetic race between detachment of the bound head and attachment of the tethered head (state 5 in Fig. 1), we can use single-molecule motility measurements to estimate 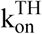. To quantify the rate of KIF1A detachment from the post-hydrolysis state, we used the ADP state as a proxy for this weakly-bound state and measured single-molecule binding durations in varying ADP concentrations (Fig. 6A and B). Microtubule off-rates at each [ADP] were obtained by fitting to the exponential dwell time distributions (Fig. 6C). A hyperbolic fit (see Methods) revealed a maximum off-rate of 0.27 ± 0.11 s^−1^ in ADP, an off-rate in the apo state of 0.09 ± 0.002 s^−1^, and a K_0.5_, representing the K_D_ of KIF1A for ADP when bound to the microtubule, of 93 ± 204 μM (Fig. 6D, fit ± 95% CI). This KIF1A off-rate in ADP is approximately 5-fold slower than for kinesin-1 and almost 7-fold slower than for kinesin-2 (Fig. 6D).^27,28,34,42^ Importantly, this KIF1A off-rate in ADP is very similar to the off-rate of the motor during a processive run, which we calculated as 0.28 ± 0.09 s^−1^ (Fig. 2D). If the detachment rate during stepping is considered simply as the off-rate in the weakly-bound state multiplied by the fraction of time in the weakly-bound state, then it follows that the motor must spend the majority of its cycle in the weakly-bound post-hydrolysis state. This implies that tethered head attachment is rate limiting, rather than hydrolysis.

**Figure 6.**
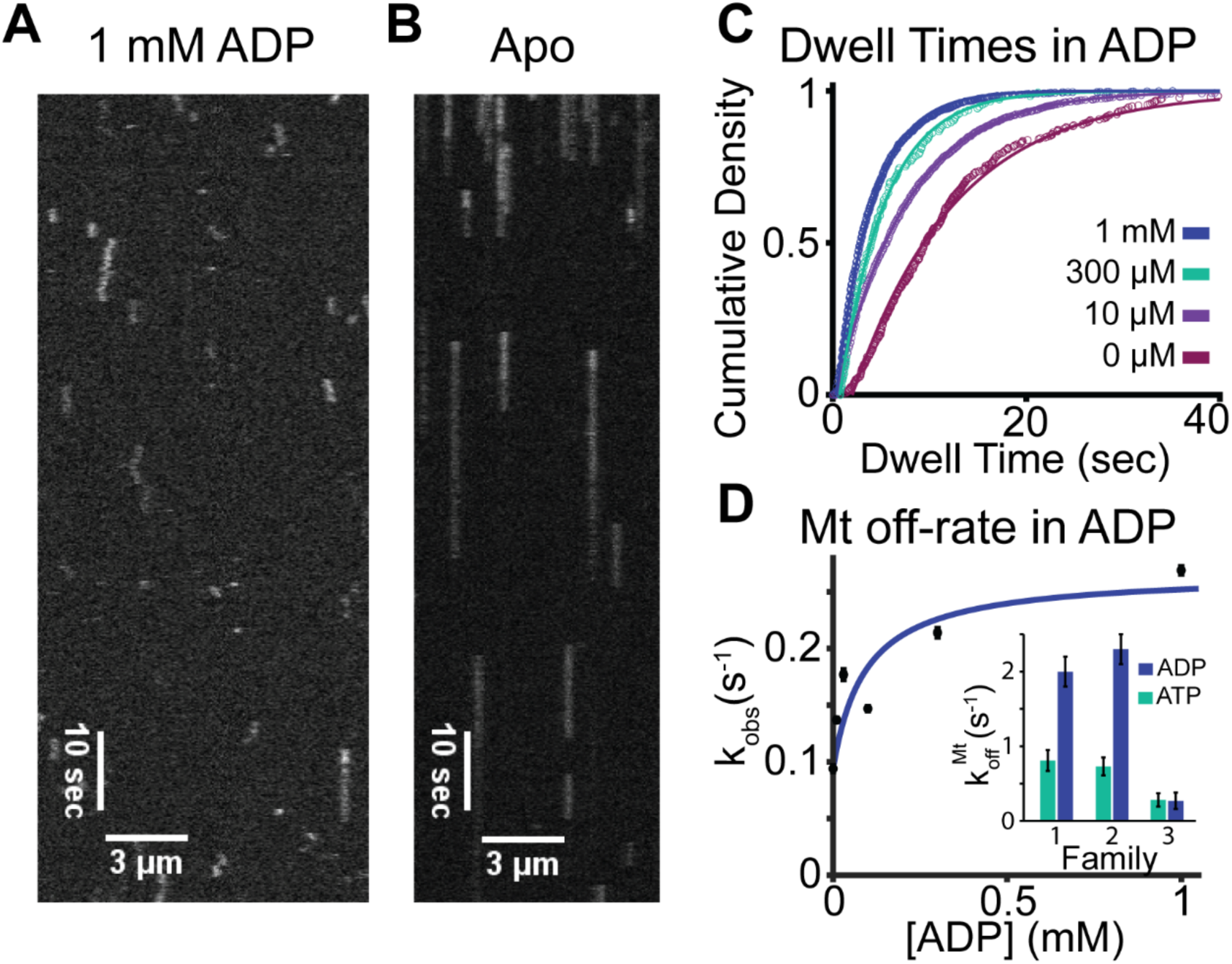
Microtubule affinity of KIF1A at varying ADP concentrations by single-molecule assay. **A**, Kymograph for KIF1A-560-GFP in 1 mM ADP at 5 fps. **B**, Kymograph for KIF1A-560-GFP in the absence of nucleotide (apo) at 5 fps. **C**, Cumulative density fit to dwell time distributions in 0, 0.01, 0.3 and 1 mM ADP gives 10.6 ± 0.2 s, 7.3± 0.02 s, 4.7 ± 0.06 s, and 3.7 ± 0.03 s, respectively. Inverse of these durations give microtubule off-rates of 0.09 ± 0.002 s^−1^, 0.14 ± 0.0002 s^−1^, 0.21 ± 0.003 s^−1^, 0.27 ± 0.002 s^−1^, respectively. Values presented as fit ± 95% confidence intervals. **D**, Microtubule off-rate of KIF1A versus the ADP concentration. Fit with Eq. 1 (See Methods) gives a maximum off-rate of 0.27 ± 0.11 s^−1^ in saturating ADP, an apo state off-rate of 0.09 ± 0.2 μM, and a K_0.5_ of 93 ± 204 μM (all fit ± 95% confidence). **Inset**, Comparing 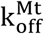 in ATP (green bars) and ADP (blue bars) in BRB80 for Kinesin-1, -2, and -3.^28^

To calculate the tethered head attachment rate more quantitatively, we can compare the motor dissociation rate in ADP to the probability the motor will detach per step it takes along the microtubule in ATP. Following ATP hydrolysis (state 5 in Fig. 1), we consider processivity as a race between the tethered head completing the forward step with a rate 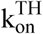and the bound head dissociating from the microtubule at a rate 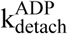.^28^ The probability of the motor detaching per step is:

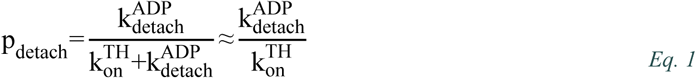

Where, for a highly processive motor, 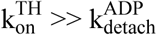. We can rearrange this kinetic race equation (Eq. 1) to solve for 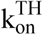. KIF1A has an estimated run length of 5.6 ± 0.4 μm (Fig 2E), meaning it takes approximately 700 steps before dissociating; thus, the probability of detaching per step is 1/700. Also, the KIF1A off-rate in ADP is 0.27 ± 0.11 s^−1^ (Fig 6D). Together, these values indicate a tethered head attachment rate of 189 ± 78 s^−1^. This rate corresponds to a duration in the 1HB state following ATP hydrolysis of 5.3 ± 2.2 ms, which is comparable to the total cycle duration of 4.5 ± 1.0 ms (Fig 2D). In support of this rate determination, we can calculate 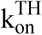 using the property that the total stepping rate is made up of the sequential transitions that make up the chemomechanical cycle (see Methods Eq. 3). Inputting the measured ATP on-rate and ADP off-rate and assuming that trailing head detachment and ATP hydrolysis are both very fast results in an estimated 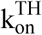 of 286 s^−1^, similar to the 189 ± 78 s^−1^ determined based on the processivity and to the 220 s^−1^ total stepping rate. To summarize, comparison of the KIF1A off-rate in the weak-binding state to either the motor off-rate in ATP or to the probability of detaching per step yields a consistent conclusion that tethered-head attachment is the rate-limiting step in the KIF1A chemomechanical cycle and that KIF1A spends the bulk of its cycle in a weak-binding 1HB state.

## DISCUSSION

In this work, we find that the KIF1A chemomechanical cycle follows the same sequence of states as established for kinesin-1 and kinesin-2,^27,34,38^ and that the motor’s fast stepping rate and superprocessivity result from differences in specific transition rates in the chemomechanical cycle. Compared to transport motors in the kinesin-1 and -2 families, the KIF1A chemomechanical cycle is distinctive in having: 1) an order of magnitude faster rear-head detachment rate; 2) a rate-limiting tethered-head attachment rate; and 3) relatively slow dissociation from the low affinity post-hydrolysis state. The measured KIF1A rate constants are summarized in Table 1. A comparison between the chemomechanical cycles of KIF1A and kinesin-1 and -2 are presented in Fig. 7 and summarized in Table 2. Below, we account for the specific motor characteristics of KIF1A in terms of our measured kinetic rates and affinities.

**Figure 7.**
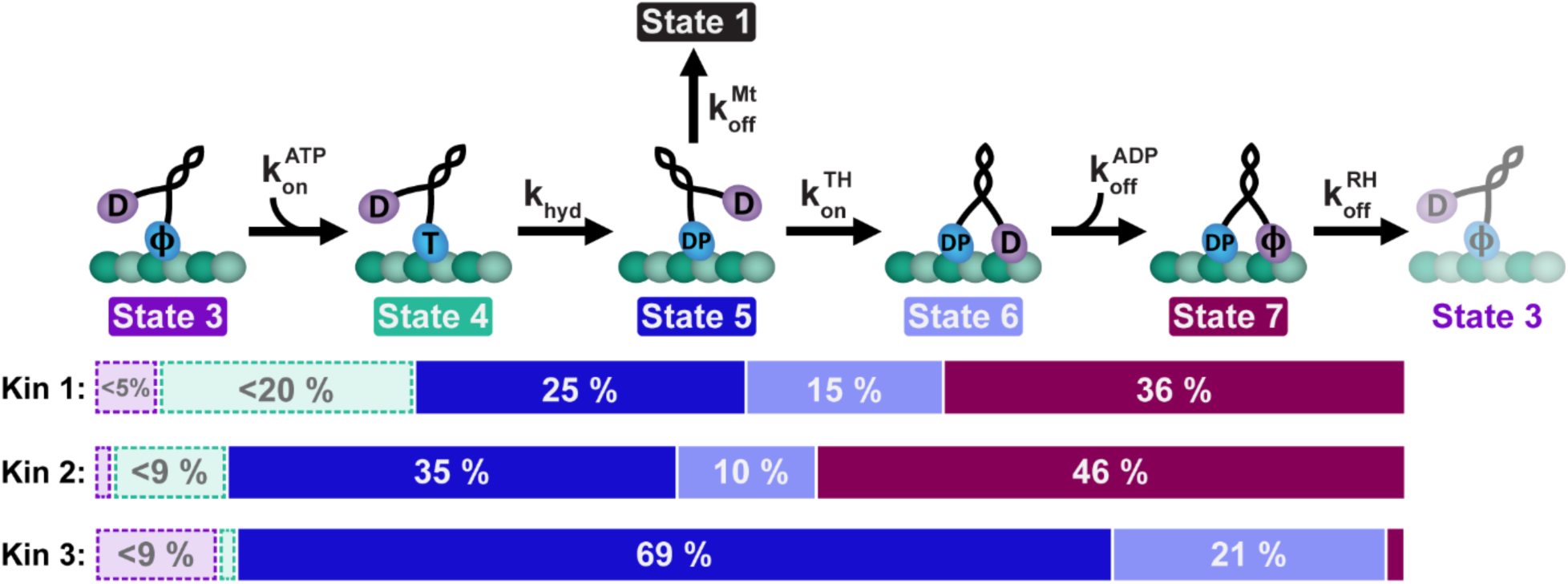
Comparison of stepping cycles for Kinesin-1, -2 and -3. The percent of time spent in each chemomechanical state is compared for Kinesin-1 (KHC),^27,28,42^ Kinesin-2 (KIF3A),^28,34^ and Kinesin-3 (KIF1A). The state numbers correspond to those in Fig. 1. Dashed boxes represent uncertainty of the duration due to experimental limitations of the ATP on-rate and ATP hydrolysis rate determinations. Small boxes without labels represent a state duration that is <1% of the cycle time. The total cycle duration used for the determination of the percentages presented here is the sum of the state durations. See Table 2 for exact values. D=ADP; T=ATP; DP= ADP-P_i;_ ϕ=Apo.

### Origin of fast Velocity

The KIF1A property that most contributes to its faster stepping rate is the rapid rear-head detachment rate. Nucleotide-triggered half-site release assays provide a convenient estimation of this transition rate because the measurement includes every transition in the chemomechanical cycle except rear-head detachment. Comparison to the overall stepping rate, which includes all transitions in the cycle, thus yields the rear-head detachment rate. For KIF1A, the ATP-triggered half-site release rate agrees with the stepping rate to within experimental error (Fig. 3C), indicating that rear head detachment is faster than we are able to measure. As a comparison, a recent kinesin-1 study measured a stepping rate of 65 s^−1^ (15.4 ms) and an ATP-triggered half-site release rate of 112 s^−1^ (8.9 ms). This yields a calculated rear-head detachment rate for kinesin-1 of 155 s^−1^ (6.5 ms), which approaches half of the overall cycle time (Fig. 7, Table 2).^27,28^ Similarly, in the slow moving kinesin-5, rear-head detachment is the rate limiting state, ensuring the motor spends the bulk of its cycle in a two-heads-bound state.^46^

The ability to quickly detach the rear-head from the microtubule appears to be in conflict with the slow microtubule off-rate of KIF1A in the ADP state, but upon closer inspection, these rates can be reconciled. It has been clearly established that this relatively high microtubule affinity of KIF1A in the ADP state results from electrostatic interaction of the positively charged loop 12 with the negatively charged C-terminal tail of tubulin.^18,24^ Additionally, the diffusive behavior of KIF1A along microtubules in ADP indicates that electrostatic interactions with any given tubulin are fleeting, and that the motor remains bound to the microtubule by renewing electrostatic interactions with different tubulin subunits along the lattice.^15,24^ Thus, the measured off-rate of 0.27 s^−1^ (Fig. 6D) in ADP does not represent the off-rate from individual tubulin but rather from the entire microtubule. Secondly, the rear-head detachment rate is thought to be accelerated by inter-head tension when the motor is in the two-heads-bound state,^47,48^ which contrasts with the unloaded off-rate in ADP. Of note, we found that the microtubule off-rate in the strong-binding apo state is more than an order of magnitude faster in KIF1A than in kinesin-1 (Fig. 6D).^49^ Thus, one possible interpretation is that in weak-binding states KIF1A is stabilized by more electrostatic interactions with the microtubule than is kinesin-1, but kinesin-1 forms greater stabilizing interactions with the microtubule in strong-binding states. Previous CryoEM and Molecular Dynamics studies have noted differences between the microtubule binding interfaces of kinesin-3 and kinesin-1,^50,51^ but they are unable to clearly account for this lower affinity in the apo state.

The faster stepping rate of KIF1A results from not only a faster rear head detachment rate, but also a faster tethered head binding rate compared to kinesin-1 and -2 (Fig. 7, Table 2).^28,34^ This faster tethered head binding rate is qualitatively consistent with the fast microtubule on-rate of KIF1A, measured by stopped flow here and from landing rates in previous single-molecule investigations.^15^ However, compared to kinesin-1, KIF1A has a 15-fold faster 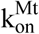, but less than two-fold faster tethered head attachment rate. Thus, the electrostatic interactions that likely determine the fast 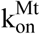, are not the dominant factor in tethered head binding during motor stepping. One potential explanation for this kinetic discrepancy is that the tethered head attachment rate is determined not by the association kinetics between the tethered head and the microtubule, but rather by the kinetics of neck linker docking. A recent structural and Molecular Dynamics study found that, compared to kinesin-1, neck linker docking in KIF1A is stabilized by fewer hydrogen bonds between the neck linker, cover strand, and catalytic core.^52^ This reduced stabilization could manifest as a slower rate of neck linker docking in KIF1A. Consistent with this, we observed a relatively slow half-site release rate in ATPγS and AMPPNP compared to kinesin-1 and -2 (Fig. 6C).^27,37,41^ In kinesin-1, AMPPNP triggers half-site release at roughly one-third the rate of ATP, consistent with ATP binding alone inducing at least partial neck linker docking.^41^ In contrast, AMPPNP-triggered half-site release in KIF1A is more than two orders of magnitude slower than ATP, which is difficult to reconcile with any degree of neck linker docking preceding ATP hydrolysis.

Our conclusion that tethered head attachment is rate-limiting for KIF1A is supported by two lines of evidence, but there are caveats. The key finding is that the off-rate in ADP is quite slow. The agreement with the motor off-rate during processive stepping means that the motor must spend the majority of its cycle in this low affinity state, and modeling processivity as a kinetic race yields a tethered-head on-rate similar to the overall stepping rate. One caveat is that we are using the motor off-rate in ADP as a model of the post-hydrolysis state. Whether the head dissociates in the ADP-Pi state and rapidly releases Pi, or whether Pi release precedes dissociation is not known. There is evidence from kinesin-1 that the ADP-Pi state is higher affinity than the ADP state.^36^ If this is the case for KIF1A, this would provide a quandary because the tethered-head on-rate would need to be slower than the overall stepping rate to explain the processivity of KIF1A. It has been suggested based on crystal structures in solution that the ADP-Pi state of KIF1A may have a lower microtubule affinity than the ADP state.^26^ However, the relevance of these structures to microtubule-docked structures is questionable and there are no supporting functional data. A second caveat is that, if tethered head attachment is rate-limiting, then it implies a very fast ATP hydrolysis rate. Hydrolysis rates for other kinesins have been indirectly estimated to be a few hundred per second (Fig. 7, Table 2),^28,34^ but the rate of hydrolysis is very difficult to measure quantitatively and is arguably the most poorly defined rate constant in the kinesin chemomechanical cycle. Nonetheless, a hydrolysis rate over 1000 s^−1^ seems unlikely, and because the KIF1A stepping cycle is so fast, rates below this imply that the time for hydrolysis is a non-negligible fraction of the cycle. In summary, our data support tethered head attachment as the sole rate limiting step, but there are caveats and a more precise estimate of this rate constant will require high-resolution head-tracking experiments as have been carried out for kinesin-1.^27,53^

### Origin of Superprocessivity and Load Sensitivity

The finding that rear-head detachment is fast and tethered head attachment is rate-limiting means that the motor spends most of its cycle in a one-head-bound state, a property that would generally be expected to reduce processivity. The key characteristic of KIF1A that determines its superprocessivity is its slow off-rate in the post-hydrolysis state (state 5 → 1, Fig. 1 and 7). This trait was observed first in the finding that an engineered KIF1A monomer in low ionic strength buffer is capable of processive transport.^24^ This electrostatic tethering thus contributes to both high velocity (by allowing fast rear head detachment) and superprocessivity (by minimizing probability of detachment during a step). However, a negative byproduct of the motor spending most of its time in a 1HB weak-binding state is that KIF1A tends to detach against applied loads.^17,29–31^ In an optical trapping assay using the *C. elegans* KIF1A, Unc104, a 1 pN applied load led to a 10-fold increase in the motor detachment rate.^19^ This effect is also seen in mixed motor assays, where minor fractions of the slower kinesin-1 mixed with the fast kinesin-3 lead to mixed motor speeds very similar to kinesin-1,^31^ and in engineered pairs of kinesin-1 and kinesin-3, where the speed of the pair is very close to the speed of kinesin-1 alone.^29^ These multi-motor assays suggest that when the slower kinesin-1 pulls against the faster kinesin-3, the kinesin-3 motors detach.

### Conclusions

Defining the KIF1A chemomechanical cycle is important both for understanding the motor’s diverse transport functions in cells and understanding how kinesins have evolved to achieve diverse mechanochemistry. From a design perspective, fast speed and superprocessivity provide competing constraints because each head must cyclically detach from the microtubule, while the dimeric motor remains associated over hundreds of steps. KIF1A does this by maximizing the rear head detachment rate and maintaining electrostatic association with the microtubule even in the weak binding post-hydrolysis state. As a result, however, the motor is sensitive to load. It may be that these motor properties have evolved for multi-motor transport where each motor feels only a small fraction of the load or where the rapid motor reattachment of KIF1A ensures a stable population of motors bound to the microtubule. The mitotic kinesin-5 motor, Eg5, provides a contrast to KIF1A in that it moves roughly 20-fold slower, is much less processive,^54^ and is able to generate large forces as teams because it spends most of its hydrolysis cycle in a two-head-bound state.^46,55,56^ Thus, by tuning their chemomechanical cycles, kinesins are able to achieve diverse mechanochemistry and carry out diverse cellular functions.

## METHODS

### Protein Constructs, Purification, and Activity Quantification

The KIF1A construct used in the biochemical assays (KIF1A-406) consisted of the motor head and neck linker domains (1-368) of *Rattus norvegicus* KIF1A followed by 61 residues (445-405) from the neck-coil domain of *Drosophila melanogaster* KHC. The KIF1A construct used for the single-molecule experiments (KIF1A-560-GFP) includes an additional 216 residues from the coiled-coil domain of *Dm*KHC followed by a C-terminal GFP. Both constructs included a C-terminal 6xHis-tag. These constructs match similar kinesin-1, -2, -5, and -7 constructs analyzed in previous studies.^34,46,48^ The bacterial expression of KIF1A-560-GFP was carried out in a 2 L flask in-house followed by Ni gravity column chromatography purification with an elution buffer containing 10 μM ATP and DTT, following published protocols.^57,58^ The elution was exchanged into storage buffer (BRB80, 10 uM ATP, 5 mM βME, 5% glycerol) and then flash frozen and stored at -80°C. The concentration of KIF1A-560-GFP was quantified using GFP absorption at 488 nm.

The KIF1A-406 construct used for biochemical experiments was bacterially expressed in a Sartorius Biostat Cplus 30 L vessel at the CSL Behring Fermentation Facility at the Pennsylvania State University. The motor was purified by Ni column chromatography on an AKTA Pure FPLC system with an elution buffer containing 10 μM ATP and DTT, following published protocols.^34,57^ Following purification, KIF1A-406 was incubated in 200 μM mADP and then buffer exchanged into BRB80 buffer (80 mM PIPES, 1 mM EGTA, 1 mM MgCl_2_, pH 6.9) plus 0.5 μM or 10 μM mADP using a PD10 G25 desalting column. Sucrose was then added to the peak fractions and aliquots flash frozen and stored at −80°C. To quantify the active motor dimer concentrations for stopped flow assays, a motor sample was incubated with 1 mM ATP to chase off the bound mADP, the fluorescence of mADP (356-nm excitation/450-nm emission) measured and converted to [mADP] using a calibration curve, the solution mADP subtracted, and the value divided by two.^34^ We found that the nearly μM mADP affinity for KIF1A in solution and competition with free ATP from the purification procedure led to underestimates of the true active motor concentration by this method. Therefore, for ATPase assays where the active concentration was critical, the active motor concentration was determined by pelleting motors in the presence of microtubules and AMPPNP, quantifying the fraction of motors remaining in the supernatant via SDS-PAGE and ImageJ gel band intensity analysis, and multiplying this relative activity by the total motor concentration determined by A_280_.

### Single-Molecule Fluorescence Tracking

Single-molecule tracking of GFP-labeled KIF1A-560 was performed on a Nikon TE2000 TIRF microscope at 25°C, as described previously.^27,47,48^ Flow cells were functionalized by flowing in 0.5 mg/ml casein, followed by full-length rigor kinesin.^27^ Taxol-stabilized microtubules, polymerized from a 1:20 ratio of Cy5-labeled (GE Healthcare) and unlabeled tubulin, were then introduced, and after a 5 min incubation, motors were introduced and imaged. KIF1A motile events were recorded at 5 or 10 fps and manually analyzed using the Kymograph Evaluation tool in FIESTA software^59^ to determine the run length, velocity and dwell times. In the ADP dwell time assays, some trials contain hexokinase to reduce the amount of ATP contamination in solution. Calculations of observed motor off-rates per concentration were done using the relation 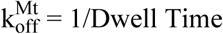.^34^ Plots of the observed off-rate as a function of ADP concentration were fit with the following equation to determine the maximum off-rate and dissociation constant of the motor for the microtubule in the nucleotide state.

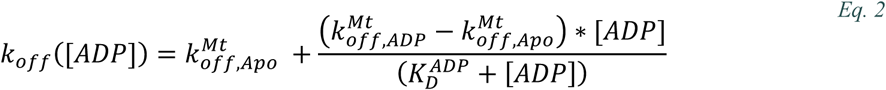

### ATPase Assays

KIF1A ATPase rates were measured by quantifying the rate of NADH conversion in an enzyme coupled reaction at varying [Mt], as described by Huang and Hackney.^34,60^ The reaction contained BRB80 with 1 mM Mg-ATP, 2 mM phosphoenolpyruvate, 1 mM MgCl_2_, 0.2 mg/ml casein, 10 μM Taxol, 0.25 mM NADH, and [1.5/100] volume of PK/LDH (Sigma P-0294). Absorbance of NADH at 340 nm over time was measured on a Molecular Devices FlexStation 3 Multi-mode Microplate Reader, converted to an ATPase rate, and divided by the active motor concentration to give the total hydrolysis cycle rate at 25°C.

### Stopped Flow Setup

Stopped-flow experiments were carried out at 25°C in BRB80 buffer using an Applied Photophysics SX20 spectrofluorometer at 356-nm excitation with an HQ480SP emission filter. Each sample trial reported is based on the fit of the average trace of 5-7 consecutive shots. Concentrations reported below are pre-mix syringe concentrations, and thus are twice the final chamber concentrations. In the Results section, all concentrations are chamber reaction concentrations.

### 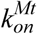 Experiments

To obtain the bimolecular on-rate for microtubule binding, 300 nM mADP exchanged KIF1A dimers in 0.5 μM free mADP were flushed against varying concentrations of taxol-stabilized microtubules in a solution of 2 mM ADP. The change in fluorescence due to release of mADP from the bound-head was fit with a double-exponential to determine the k_obs_. The fast phase of the exponential fits were plotted versus the microtubule concentration and fit linearly to obtain 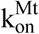. The slow phase was attributed to slower mADP release by the second head.^34^

### Half-Site Reactivity Experiment

300 nM of mADP labeled KIF1A was flushed against a solution of 2 μM taxol-stabilized microtubules with or without 2 mM ATP. The change in fluorescence due to mADP release from the bound-head(s) was fit with a single-exponential to determine the amplitude, and the relative amplitudes compared in the presence and absence of ATP.^40^

### Nucleotide-stimulated Half-site Release Assays

To establish a one-head-bound complex, 300 nM of mADP exchanged KIF1A dimers was incubated with 6 μM taxol-stabilized microtubules. This solution was then flushed against varying concentrations of ATP, ATPγS or AMPPNP. The change in fluorescence due to release of mADP from the tethered-head was fit with a single-exponential, the rates plotted against the nucleotide concentration, and the curve fit with the Michaelis-Menten equation to obtain the maximum release rate and K_0.5_. ^34,41^

### Nucleotide Exchange Experiments

To determine the ADP solution off-rate, 0.3 μM KIF1A in a solution of 0.5 μM free ADP was flushed against 10 μM mADP. In this configuration, the exponential increase in fluorescence from the binding of mADP is rate limited by the off-rate of ADP in solution. In the complementary assay to determine the mADP solution off-rate, 0.3 μM mADP-exchanged KIF1A dimers in a solution of 0.5 μM free mADP was flushed against 2 mM ADP. The exponential decrease in fluorescence was fit to obtain the off-rate of mADP in solution.

To determine the unstrained mADP exchange rate, 1 μM mADP-exchanged KIF1A dimers were combined with 5 μM taxol-stabilized microtubules and 0.5 μM mADP to achieve a one-head-bound KIF1A-Mt complex. This solution was flushed against varying concentrations of mADP and the increase in fluorescence due to mADP binding fit to an exponential. To determine the strained mADP exchange rate, 2 μM KIF1A was pre-incubated with 10 μM microtubules and 100 μM AMPPNP to obtain a two-heads-bound complex with AMPPNP in the rear head and no nucleotide in the leading head.^34,43,44,46^ This complex was flushed against varying concentrations of mADP and the rise in fluorescence due to mADP binding fit to an exponential. Due to the high free [mADP] in both of these assays, mADP binding was monitored by exciting at 280-nm and measuring the FRET signal between Trp in the motor domain and the mADP.^61^ For both the unstrained and strained exchange assays, the exponential fits began at 2 ms, due to the instrument dead time. To obtain the ADP on- and off-rates, the resulting k_obs_ were plotted versus the mADP concentration and fit linearly with the equation k_obs_= k_on_*[mADP]+k_off_.

### Calculations

State transition durations within the chemomechanical cycle were calculated using the following relationship:

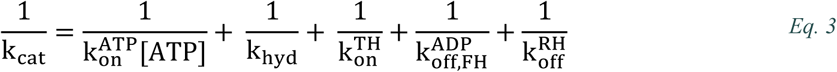

Additionally, the relationship between the total step duration and the time for half-site release is as follows:

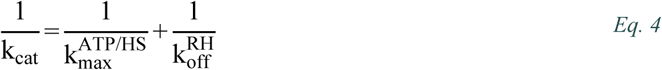

### Run Length Correction for Finite Microtubule Lengths

KIF1A has a long run length which results in a significant fraction of motors that run off the microtubule end, which if not accounted for, leads to an underestimate of the run length. Thus, motor run lengths were corrected for finite microtubule lengths, as follows.

For every event, the run length was recorded, along with whether the motor dissociated from some point along the microtubule or ran off the end. Motor stepping was assumed to be history independent and thus the run lengths were assumed to be exponentially distributed with a mean run length of θ = 1/*λ*. If the microtubule was infinitely long, the standard model for the run length would have probability density:

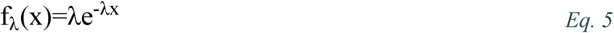

The run lengths for n motors corresponding to our observations are {X_1_, X_2_, …, X_n_}. For motors that run off the end of the microtubule, we know the distance from the landing point of the motor to the end of the microtubule and notate the value as *t*_*i*_. The measured run lengths, Y_i_, including events that run off the end, are the minimum of the true run length, X_i_, and the distance to the end of the microtubule, t_i_:

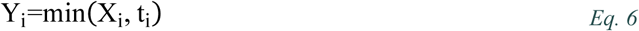

We also define a variable W_i_, denoting whether the motor dissociated normally from the lattice (W_i_ = 1) or ran off the end (W_i_ = 0). Our data will then be *Y*_*i*_, *W*_*i*_ with *t*_*i*_ serving as a known covariate. Our goal is to solve for the rate of dissociation (in inverse distance), *λ*.

The log of the likelihood function is defined as:

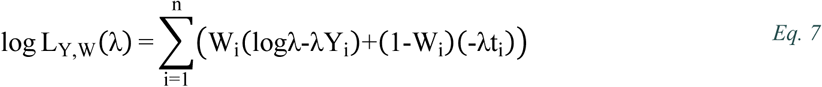

To maximize the likelihood, we take the derivative with respect to *λ* and set it to zero. Then, because we define the mean run length as θ = 1/*λ*, we can simplify to the following equation:

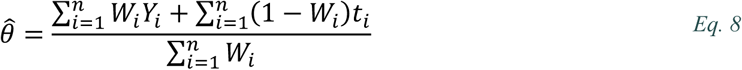

Under some broad regularity conditions, the asymptotic variance for a maximum likelihood estimator is the reciprocal of the Fisher information. So, 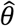 should be approximately normally distributed with mean θ and a variance of:

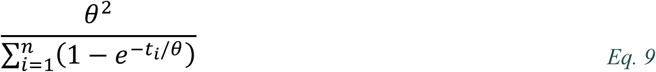

Assuming the average of the Y_i_ is defined as

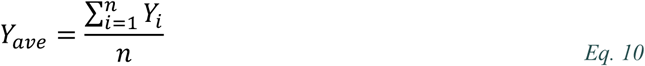

We can define t_i_ = Y_i_ and simplify the maximum likelihood estimator equation to:

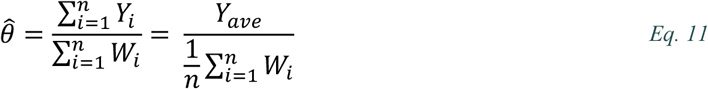

As such, the denominator represents the fraction of motors that detach normally from the lattice. Leading to the following interpretation:

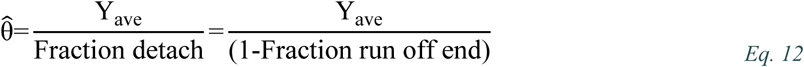

So, if all motors detach normally, then the run length is the average, but, for example, if half of the motors reach the end, then the run length is corrected up by a factor of 2. This correction should apply generally for processes that generate exponential distributions with censoring, such as photobleaching. The correction is similar to the Kaplan-Meier estimate that was used for run length corrections by Ruhnow *et al*, but has a simpler form.^62^ In addition, we are using an asymptotic result for the variance as opposed to the bootstrap method found in Ruhnow *et al*.

## Acknowledgements

The authors thank members of Hancock Lab for their contributions and helpful discussions.

This work was supported by NIH grant number R01GM076476 to W.O.H.

## REFERENCES

(1) Hirokawa, N.; Noda, Y.; Tanaka, Y.; Niwa, S. Kinesin Superfamily Motor Proteins and Intracellular Transport. Nat. Rev. Mol. Cell Biol. 2009, 10, 682–696.

(2) Hirokawa, N.; Niwa, S.; Tanaka, Y. Molecular Motors in Neurons: Transport Mechanisms and Roles in Brain Function, Development, and Disease. Neuron 2010, 68 (4), 610–638.

(3) Siddiqui, N.; Straube, A. Intracellular Cargo Transport by Kinesin-3 Motors. Biochem. 2017, 82 (7), 803–815.

(4) Okada, Y.; Yamazaki, H.; Sekine-Aizawa, Y.; Hirokawa, N. The Neuron-Specific Kinesin Superfamily Protein KIF1A Is a Uniqye Monomeric Motor for Anterograde Axonal Transport of Synaptic Vesicle Precursors. Cell 1995, 81 (5), 769–780.

(5) Hirokawa, N. Kinesin and Dynein Superfamily Proteins and the Mechanism of Organelle Transport. Science 1998, 279 (5350), 519–526.

(6) Pennings, M.; Schouten, M. I.; Gaalen, J. van; Meijer, R. P. P. P.; Bot, S. T. de; Kriek, M.; Saris, C. G. J. J.; Berg, L. H. van den; Es, M. A. van; Zuidgeest, D. M. H. H.; et al. KIF1A Variants Are a Frequent Cause of Autosomal Dominant Hereditary Spastic Paraplegia. Eur. J. Hum. Genet. 2020, 28 (1), 40–49.

(7) Yonekawa, Y.; Harada, A.; Okada, Y.; Funakoshi, T.; Kanai, Y.; Takei, Y.; Terada, S.; Noda, T.; Hirokawa, N. Defect in Synaptic Vesicle Precursor Transport and Neuronal Cell Death in KIF1A Motor Protein–Deficient Mice. J. Cell Biol. 1998, 141 (2), 431–441.

(8) Chiba, K.; Takahashi, H.; Chen, M.; Obinata, H.; Arai, S.; Hashimoto, K.; Oda, T.; McKenney, R. J.; Niwa, S. Disease-Associated Mutations Hyperactivate KIF1A Motility and Anterograde Axonal Transport of Synaptic Vesicle Precursors. Proc. Natl. Acad. Sci. 2019, 116 (37), 18429–18434.

(9) Miki, H.; Okada, Y.; Hirokawa, N. Analysis of the Kinesin Superfamily: Insights into Structure and Function. Trends Cell Biol. 2005, 15 (9), 467–476.

(10) Hirokawa, N.; Takemura, R. Molecular Motors and Mechanisms of Directional Transport in Neurons. Nat. Rev. Neurosci. 2005, 6 (3), 201–214.

(11) Miki, H.; Setou, M.; Kaneshiro, K.; Hirokawa, N. All Kinesin Superfamily Protein, KIF, Genes in Mouse and Human. Proc. Natl. Acad. Sci. 2001, 98 (13), 7004–7011.

(12) Lawrence, C. J.; Dawe, R. K.; Christie, K. R.; Cleveland, D. W.; Dawson, S. C.; Endow, S. A.; Goldstein, L. S. B.; Goodson, H. V; Hirokawa, N.; Howard, J.; et al. A Standardized Kinesin Nomenclature. J. Cell Biol. 2004, 167 (1), 19–22.

(13) Lessard, D. V.; Zinder, O. J.; Hotta, T.; Verhey, K. J.; Ohi, R.; Berger, C. L. Polyglutamylation of Tubulin’s C-Terminal Tail Controls Pausing and Motility of Kinesin-3 Family Member KIF1A. J. Biol. Chem. 2019, 294 (16), 6353–6363.

(14) Soppina, V.; Norris, S. R.; Dizaji, A. S.; Kortus, M.; Veatch, S.; Peckham, M.; Verhey, K. J. Dimerization of Mammalian Kinesin-3 Motors Results in Superprocessive Motion. PNAS 2014, 111 (15), 5562–5567.

(15) Soppina, V.; Verhey, K. J. The Family-Specific K-Loop Influences the Microtubule on-Rate but Not the Superprocessivity of Kinesin-3 Motors. Mol. Biol. Cell 2014, 25 (14), 2161–2170.

(16) Guedes-Dias, P.; Nirschl, J. J.; Abreu, N.; Tokito, M. K.; Janke, C.; Magiera, M. M.; Holzbaur, E. L. F. F. Kinesin-3 Responds to Local Microtubule Dynamics to Target Synaptic Cargo Delivery to the Presynapse. Curr. Biol. 2019, 29 (2), 268–282.e8.

(17) Okada, Y.; Higuchi, H.; Hirokawa, N. Processivity of the Single-Headed Kinesin KIF1A through Binding to Tubulin. Nature 2003, 424 (6948), 574–577.

(18) Okada, Y.; Hirokawa, N. Mechanism of the Single-Headed Processivity: Diffusional Anchoring between the K-Loop of Kinesin and the C Terminus of Tubulin. Proc. Natl. Acad. Sci. 2000, 97 (2), 640–645.

(19) Tomishige, M. Conversion of Unc104/KIF1A Kinesin into a Processive Motor After Dimerization. Science 2002, 297 (5590), 2263–2267.

(20) Al-Bassam, J.; Cui, Y.; Klopfenstein, D.; Carragher, B. O.; Vale, R. D.; Milligan, R. A. Distinct Conformations of the Kinesin Unc104 Neck Regulate a Monomer to Dimer Motor Transition. J. Cell Biol. 2003, 163 (4), 743–753.

(21) Rashid, D. J.; Bononi, J.; Tripet, B. P.; Hodges, R. S.; Pierce, D. W. Monomeric and Dimeric States Exhibited by the Kinesin-Related Motor Protein KIF1A. J. Pept. Res. 2005, 65 (6), 538–549.

(22) Kikkawa, M.; Okada, Y.; Hirokawa, N. 15 Å Resolution Model of the Monomeric Kinesin Motor, KIF1A. Cell 2000, 100 (2), 241–252.

(23) Kikkawa, M.; Sablin, E. P.; Okada, Y.; Yajima, H.; Fletterick, R. J.; Hirokawa, N. Switch-Based Mechanism of Kinesin Motors. Nature 2001, 411 (24), 439–445.

(24) Okada, Y.; Hirokawa, N. A Processive Single-Headed Motor: Kinesin Superfamily Protein KIF1A. Science 1999, 283 (5405), 1152–1157.

(25) Liu, F.; Ji, Q.; Wang, H.; Wang, J. Mechanochemical Model of the Power Stroke of the Single-Headed Motor Protein KIF1A. J. Phys. Chem. B 2018, 122 (49), 11002–11013.

(26) Nitta, R.; Kikkawa, M.; Okada, Y.; Hirokawa, N. KIF1A Alternately Uses Two Loops to Bind Microtubules. Science 2004, 305 (5684), 678–683.

(27) Mickolajczyk, K. J.; Deffenbaugh, N. C.; Ortega Arroyo, J.; Andrecka, J.; Kukura, P.; Hancock, W. O. Kinetics of Nucleotide-Dependent Structural Transitions in the Kinesin-1 Hydrolysis Cycle. Proc. Natl. Acad. Sci. 2015, 112 (52), E7186–E7193.

(28) Mickolajczyk, K. J.; Hancock, W. O. Kinesin Processivity Is Determined by a Kinetic Race from a Vulnerable One-Head-Bound State. Biophys. J. 2017, 112 (12), 2615–2623.

(29) Norris, S. R.; Soppina, V.; Dizaji, A. S.; Schimert, K. I.; Sept, D.; Cai, D.; Sivaramakrishnan, S.; Verhey, K. J. A Method for Multiprotein Assembly in Cells Reveals Independent Action of Kinesins in Complex. J. Cell Biol. 2014, 207 (3), 393–406.

(30) Arpağ, G.; Norris, S. R.; Mousavi, S. I.; Soppina, V.; Verhey, K. J.; Hancock, W. O.; Tüzel, E. Motor Dynamics Underlying Cargo Transport by Pairs of Kinesin-1 and Kinesin-3 Motors. Biophys. J. 2019, 116 (6), 1115–1126.

(31) Ker Arpa, G.; Shastry, S.; Hancock, W. O.; Tü Zel, E.; Arpağ, G.; Shastry, S.; Hancock, W. O.; Tüzel, E.; Ker Arpa, G.; Shastry, S.; et al. Transport by Populations of Fast and Slow Kinesins Uncovers Novel Family-Dependent Motor Characteristics Important for In Vivo Function. Biophys. J. 2014, 107 (8), 1896–1904.

(32) Yildiz, A.; Tomishige, M.; Vale, R. D.; Selvin, P. R. Kinesin Walks Hand-Over-Hand. Science 2004, 303 (5658), 676–678.

(33) Mickolajczyk, K. J.; Cook, A. S. I.; Jevtha, J. P.; Fricks, J.; Hancock, W. O. Insights into Kinesin-1 Stepping from Simulations and Tracking of Gold Nanoparticle-Labeled Motors. Biophys. J. 2019, 117 (2), 331–345.

(34) Chen, G.-Y.; Arginteanu, D. F. J.; Hancock, W. O. Processivity of the Kinesin-2 KIF3A Results from Rear Head Gating and Not Front Head Gating. J. Biol. Chem. 2015, 290 (16), 10274–10294.

(35) Andreasson, J. O. L.; Shastry, S.; Hancock, W. O.; Block Correspondence, S. M.; Block, S. M. The Mechanochemical Cycle of Mammalian Kinesin-2 KIF3A/B under Load. Curr. Biol. 2015, 25 (9), 1166–1175.

(36) Milic, B.; Andreasson, J. O. L.; Hancock, W. O.; Block, S. M. Kinesin Processivity Is Gated by Phosphate Release. Proc. Natl. Acad. Sci. 2014, 111 (39), 14136–14140.

(37) Andreasson, J. O.; Milic, B.; Chen, G.-Y.; Guydosh, N. R.; Hancock, W. O.; Block, S. M. Examining Kinesin Processivity within a General Gating Framework. Elife 2015, 4, e07403.

(38) Hancock, W. O. The Kinesin-1 Chemomechanical Cycle: Stepping Toward a Consensus. Biophys. J. 2016, 110 (6), 1216–1225.

(39) Vale, R. D. The Way Things Move: Looking Under the Hood of Molecular Motor Proteins. Science 2000, 288 (5463), 88–95.

(40) Hackney, D. D. Evidence for Alternating Head Catalysis by Kinesin during Microtubule-Stimulated ATP Hydrolysis. Proc. Natl. Acad. Sci. U. S. A. 1994, 91 (15), 6865–6869.

(41) Ma, Y. Z.; Taylor, E. W. Interacting Head Mechanism of Microtubule-Kinesin ATPase. J. Biol. Chem. 1997, 272 (2), 724–730.

(42) Feng, Q.; Mickolajczyk, K. J.; Chen, G.-Y.; Hancock, W. O. Motor Reattachment Kinetics Play a Dominant Role in Multimotor-Driven Cargo Transport. Biophys. J. 2018, 114 (2), 400–409.

(43) Schnapp, B. J.; Crise, B.; Sheetz, M. P.; Reese, T. S.; Khan, S. Delayed Start-up of Kinesin-Driven Microtubule Gliding Following Inhibition by Adenosine 5’-[Beta,Gamma-Imido]Triphosphate. Proc. Natl. Acad. Sci. 1990, 87 (24), 10053–10057.

(44) Guydosh, N. R.; Block, S. M. Backsteps Induced by Nucleotide Analogs Suggest the Front Head of Kinesin Is Gated by Strain. Proc. Natl. Acad. Sci. U. S. A. 2006, 103 (21), 8054–8059.

(45) Hackney, D. D. Kinesin ATPase: Rate-Limiting ADP Release. Proc. Natl. Acad. Sci. U. S. A. 1988, 85 (17), 6314–6318.

(46) Chen, G.-Y.; Mickolajczyk, K. J.; Hancock, W. O. The Kinesin-5 Chemomechanical Cycle Is Dominated by a Two-Heads-Bound State. J. Biol. Chem. 2016, 291 (39), 20283–20294.

(47) Shastry, S.; Hancock, W. O. Neck Linker Length Determines the Degree of Processivity in Kinesin-1 and Kinesin-2 Motors. Curr. Biol. 2010, 20 (10), 939–943.

(48) Shastry, S.; Hancock, W. O. Interhead Tension Determines Processivity across Diverse N-Terminal Kinesins. Proc. Natl. Acad. Sci. 2011, 108 (39), 16253–16258.

(49) Yajima, J.; Alonso, M. C.; Cross, R. A.; Toyoshima, Y. Y. Direct Long-Term Observation of Kinesin Processivity at Low Load. Curr. Biol. 2002, 12 (01), 301–306.

(50) Scarabelli, G.; Soppina, V.; Yao, X.-Q. Q.; Atherton, J.; Moores, C. A.; Verhey, K. J.; Grant, B. J. Mapping the Processivity Determinants of the Kinesin-3 Motor Domain. Biophys. J. 2015, 109 (8), 1537–1540.

(51) Atherton, J.; Farabella, I.; Yu, I. M.; Rosenfeld, S. S.; Houdusse, A.; Topf, M.; Moores, C. A. Conserved Mechanisms of Microtubule-Stimulated ADP Release, ATP Binding, and Force Generation in Transport Kinesins. Elife 2014, 3, e03680.

(52) Ren, J.; Zhang, Y.; Wang, S.; Huo, L.; Lou, J.; Feng, W. Structural Delineation of the Neck Linker of Kinesin-3 for Processive Movement. J. Mol. Biol. 2018, 430 (14), 2030–2041.

(53) Isojima, H.; Iino, R.; Niitani, Y.; Noji, H.; Tomishige, M. Direct Observation of Intermediate States during the Stepping Motion of Kinesin-1. Nat. Chem. Biol. 2016, 12 (4), 290–297.

(54) Valentine, M. T.; Fordyce, P. M.; Krzysiak, T. C.; Gilbert, S. P.; Block, S. M. Individual Dimers of the Mitotic Kinesin Motor Eg5 Step Processively and Support Substantial Loads in Vitro. Nat. Cell Biol. 2006, 8 (5), 470–476.

(55) Saunders, A. M.; Powers, J.; Strome, S.; Saxton, W. M. Kinesin-5 Acts as a Brake in Anaphase Spindle Elongation. Curr. Biol. 2007, 17 (12), 453–454.

(56) Shimamoto, Y.; Forth, S.; Kapoor, T. M. Measuring Pushing and Braking Forces Generated by Ensembles of Kinesin-5 Crosslinking Two Microtubules. Dev. Cell 2015, 34 (6), 669–681.

(57) Uppalapati, M.; Huang, Y.; Shastry, S.; Jackson, T. N.; Hancock, W. O. Microtubule Motors in Microfluidics. Methods Bioeng. Microfabr. Microfluid. 2009, 311–337.

(58) Gicking, A. M.; Wang, P.; Liu, C.; Mickolajczyk, K. J.; Guo, L.; Hancock, W. O.; Qiu, W. The Orphan Kinesin PAKRP2 Achieves Processive Motility via a Noncanonical Stepping Mechanism. Biophys. J. 2019, 116 (7), 1270–1281.

(59) Ruhnow, F.; Zwicker, D.; Diez, S. Tracking Single Particles and Elongated Filaments with Nanometer Precision. Biophys. J. 2011, 100 (11), 2820–2828.

(60) Huang, T.-G. G.; Suhan, J.; Hackney, D. D.; Suhan, J.; Hackney, D. D.; Suhan, J.; Hackney, D. D. Drosophila Kinesin Motor Domain Extending to Amino Acid Position 392 Is Dimeric When Expressed in Escherichia Coli. J. Biol. Chem. 1994, 269 (23), 16502–16507.

(61) Cheng, J. Q.; Jiang, W.; Hackney, D. D. Interaction of Mant-Adenosine Nucleotides and Magnesium with Kinesin. Biochemistry 1998, 37 (15), 5288–5295.

(62) Ruhnow, F.; Kloβ, L.; Diez, S. Challenges in Estimating the Motility Parameters of Single Processive Motor Proteins. Biophys. J. 2017, 113 (11), 2433–2443.

